# Testing the possibility of model-based Pavlovian control of attention to threat

**DOI:** 10.1101/288027

**Authors:** D Talmi, M Slapkova, MJ Wieser

## Abstract

Signals for reward or punishment attract attention preferentially, a principle termed ‘value-modulated attention capture’ (VMAC). The mechanisms that govern the allocation of attention resources can be described with a terminology that is more often applied to the control of overt behaviours, namely, the distinction between instrumental and Pavlovian control, and between model-free and model-based control. While instrumental control of VMAC can be either model-free or model-based, it is not known whether Pavlovian control of VMAC can be model-based. To decide whether this is possible we measured Steady-State Visual Evoked Potentials (SSVEPs) while 20 healthy adults took part in a novel task. During the learning stage participants underwent aversive threat conditioning with two CSs, one that predicted pain (CS+) and one that predicted safety (CS−). Instructions given prior to the test stage in the task allowed participants to infer whether novel, ambiguous CSs (new_CS+/ new_CS−) were threatening or safe. Correct inference required combining stored internal representations and new propositional information, the hallmark of model-based control. SSVEP amplitudes quantified the amount of attention allocated to novel CSs on their very first presentation, before they were ever reinforced. We found that SSVEPs were higher for new_CS+ than new_CS−. This result is potentially indicative of model-based Pavlovian control of VMAC, but additional controls are necessary to verify this conclusively. This result underlines the potential transformative role of information and inference in emotion regulation.

## Introduction

Regulating established emotional responses upon receipt of novel information can be adaptive. For example, it would be advantageous if, when a patient is told that their new bottle of medication has fewer side effects than the one they used for some years prior, they immediately down-regulated their feeling of anxiety about administering this medication, as well as concomitant cognitive, physiological and behavioural responses, such as increased attention to the bottle. In this example the bottle of medication has become, through the years of being paired with unpleasant side effects, a signal for aversive outcomes. Signals for reward or punishment are known to attract attention preferentially, a principle termed Value-Modulated Attention Capture (VMAC, Le Pelley et al., 2016).

The mechanisms that govern attention allocation can be productively described with a terminology more commonly applied to the control of overt behaviours – the distinction between instrumental/Pavlovian control, and model-free/model-based control (Dayan & Berridge, 2014). The distinction between instrumental and Pavlovian control has to do with the dependencies between behaviour and outcome (Mackintosh, 1983). In an instrumental learning task outcomes depend on agents’ behaviour, so agents act in order to increase utility – either increase the likelihood of reward, or decrease the likelihood of punishment. An example for an instrumental control of VMAC is increased attention to stimuli when told that doing so will be remunerated. By contrast, Pavlovian control refers to behaviour that is triggered by stimuli that predict reward or punishment, even when the outcomes are independent of the agent’s behaviour, such as freezing in response to a tone that predicts a pain. While Pavlovian control of VMAC was established in the case of rewarding stimuli, there are also demonstrations of the same effect with aversive stimuli (Van Damme, Crombez, Hermans, Koster, & Eccleston, 2006; L. Wang, Yu, & Zhou, 2013; Wentura, Müller, & Rothermund, 2014). One of the surest ways to be convinced that a particular behaviour is controlled by a Pavlovian, rather than an instrumental, process is when it incurs a loss (Dayan, Niv, Seymour, & Daw, 2006). Pavlovian control of VMAC was elegantly demonstrated in an experiment that used an omission schedule, where attending distractors that signalled reward magnitude resulted in the omission of the reward (Le Pelley, Pearson, Griffiths, & Beesley, 2015). Because increased attention to distractors that predicted high (compared to low) reward decreased the likelihood of reward and was never itself rewarded, VMAC in that experiment could not be attributed to instrumental control. Instead, the findings revealed a Pavlovian control of VMAC. Subsequent work showed that these effects extended to separate tasks (Bucker & Theeuwes, 2017).

The distinction between instrumental and Pavlovian control is orthogonal to the distinction between model-based and model-free control (Daw, Niv, & Dayan, 2005; Dayan & Berridge, 2014). A model, according to Dayan and Berridge, is an internal representation of stimuli, situations and environmental circumstances, which supports prospective cognition. Model-based responses are executed when we infer, based on our model of the environment, that responding in a particular way would maximise our expected utility. Model-based control can be contrasted to model-free control, which depends on the accumulated average experience agents have with the outcomes associated with a particular environmental state. Model-based control allows us to respond flexibly to a volatile, changing environment; model-free control gives us the wisdom of the average experience. The example above for instrumental control, where participants attend stimuli when told they will be rewarded for doing so, is likely to be an example for model-based instrumental control. This is because propositional information in instructions shape our model of the environment; we can take up a suggestion or follow an instruction regardless of previous experience in a task (Olsson & Phelps, 2004). Model-based instrumental responses, such as those informed by instructions, can become model-free if they are repeatedly executed and reinforced (Yin & Knowlton, 2006). For example, with repeated pairing between attention to certain stimuli and reward attainment participants acquire a habit to attend to those stimuli. The model-free nature of this behaviour is demonstrated when participants continue to pay preferential attention to these stimuli even when further reinforcement is unlikely (Luque et al., 2017).

Here we ask whether model-based Pavlovian control of VMAC is possible. Previous experiments in animals have demonstrated model-based Pavlovian control of overt behaviour. For example, placing animals in entirely new states, such as a salt-deprived state, instantly transforms the learned aversive value of a lever that predicts a salty taste (Robinson & Berridge, 2013). The opening example demonstrates what Model-based Pavlovian control of VMAC might looks like in humans: the information on the new medication revises the patient’s model of the environment, yielding new inferences that instantaneously transform the value they assign to the bottle and, consequently, the attention she pays to it. Not much is currently known about Model-based Pavlovian control in humans, although a recent study suggested that a model-based algorithm fitted conditioned threat response in the amygdala better than model-free algorithms (Prévost, McNamee, Jessup, Bossaerts, & O’Doherty, 2013), and the distinction between model-based and model-free Pavlovian control resembles the one between cognitive and emotional control of Pavlovian responses in aversive conditioning tasks (Sevenster, Beckers, & Kindt, 2012).

It is not, at present, known whether model-based Pavlovian control of VMAC is possible. Because Pavlovian control of VMAC was evident even when participants had plenty of opportunity to learn that the way they were allocating their attention was detrimental, and even when they were fully informed about the nature of the omission schedule (Pearson, Donkin, Tran, Most, & Le Pelley, 2015), the Pavlovian control of VMAC may always be model-free (Le Pelley et al., 2016). The same conclusion appears to be supported by findings that instructed extinction did not modulate the classically-conditioned potentiated startle responses (Sevenster et al., 2012). Our aim was to test this contention by using an optimised task. Evidence for model-based, Pavlovian control of VMAC will confirm, in the domain of internal resource allocation, the distinction between model-free and model-based Pavlovian control of incentive behaviour.

In the conditioning stage of the task participants passively viewed two Conditioned Stimuli (CSs), which fully predicted a painful electric shock (CS+) or the absence of shock (CS−). We measured the Steady-State Visual Evoked Potential (SSVEP), a validated neural signal of visual attention (Matthias M. Müller, Teder-Sälejärvi, & Hillyard, 1998; Norcia, Appelbaum, Ales, Cottereau, & Rossion, 2015). The SSVEP is known to be sensitive to VMAC, specifically value-modulated attentional engagement and disengagement processes that occur from 500ms after stimulus presentation (Miskovic & Keil, 2013; Wieser, Miskovic, & Keil, 2016), so we expected greater SSVEP amplitudes for the CS+ than the CS−. The test stage included a single presentation of two ambiguous, novel CSs, which we refer to as “New_CSs” to contrast them to the “old_CSs” that participants experienced during the conditioning stage. New_CSs were constructed such that their value could not be predicted by previous experience. Before the test, participants received propositional information that, when combined with their learned internal representation of the CSs, allowed them to infer the prospective value of new_CSs. We only measured attention to the very first presentation of the new_CSs, before they were ever reinforced (the entire task was repeated several times, but new stimuli were used in each repetition).

While not a formal test of the model-based or the Pavlovian nature of this form of control over VMAC, we argue that it would be difficult to account for differential attention to new_CSs in any other way, an issue we return to in the discussion section. We hypothesised that participants would be able to utilise stored information jointly with propositional information to control attention allocation, and therefore expected greater SSVEP amplitudes for the new_CS+ compared to the new_CS−.

## Materials and methods

### Participants

Twenty seven undergraduate students from the University of Manchester participated in the study in exchange for course credits. None of the participants reported personal or family history of photic epilepsy, none were taking centrally-acting medication, none had a history of psychiatric or neurological disorders, and all had normal or corrected-to-normal vision. The experiment was approved by the University of Manchester ethics committee. Three participants did not complete the study and one participant did not exhibit an SSVEP signal. Three participants were excluded because they failed the contingency awareness criterion (see below). This resulted in a total of 20 participants (6 males, mean age 19.5, SD=1.15).

### Apparatus

The amplitude of the electric skin stimulation which served as a US (see below) was controlled by a Digitimer DS5, an Isolated Bipolar Constant Current Stimulator (Digitimer DS5 2000, Digitimer Ltd., Welwyn Garden City, UK). The DS5 produces an isolated constant current stimulus proportional to a voltage applied at its input. For reasons of participant safety this stimulator is limited to delivering a maximum of 10V/10mA. Participants received the stimulation through a ring electrode built in-house (Medical Physics, Salford Royal Hospital) attached to the DS5. To ensure adequate conductance between the electrode and the skin, the back of each participant’s hand was prepared with Nuprep Skin Preparation Gel and Ten20 Conductive Paste prior to attaching the electrode. Transcutaneous electrical stimulation activates myelinated Aβ somatosensory fibres as well as Aδ nociceptive fibres (Hird, Jones, Talmi, & El-Deredy, 2018) and can therefore cause both a sensation of touch and a sensation of pain.

The experiment was implemented using the Psych toolbox on a Matlab (The Mathworks Inc., Natick, MA, USA) platform. The voltage inputs to the DS5 were sent from Matlab through a data acquisition interface (National Instruments, Austin, TX, USA). The behavioural ratings were taken on Microsoft Excel.

### Materials

#### CSs

Stimuli resembled Navon figures (Navon, 1977), in that they were composed of global and local shapes where the outline of the large ‘global’ shape was created out of smaller ‘local’ shapes. To create these stimuli we first created 48 unique shapes using Adobe Illustrator, each with a black outline and white filling. These shapes were divided into 24 pairs so that the two shapes in each pair were visually dissimilar (e.g. an arrow and a star). Each pair was used to create a subset of 4 Navon figures, as shown in Figure 1. Two were congruent (e.g. a global arrow made of local arrows, a global star made of local stars), and two incongruent (e.g. a global arrow made of local stars, a global star made of local arrows). In total, the experiment used 24 such four-figure subsets (96 Navon figures). All figures were created and presented in grayscale to minimise differences in colours and luminance. Four-figure subsets were randomly allocated to experimental block. The two congruent figures were randomly allocated to the “old_CS+” and “old_CS−“ conditions. There were three types of task blocks, as described below, termed global, local, and control blocks. The new_CS+ in ‘Global’ blocks used the global attribute of the old_CS+ and the local attribute of the old_CS−. The new_CS+ in ‘Local’ blocks used the global attribute of the old_CS− and the local attribute of the old_CS+. Two additional four-figure subsets were used for the 2 practice blocks. The figures in practice blocks were created from 4 letters with one four-figure subset consisting of ‘H’ and ‘O’, and the other one ‘Z’ and ‘I’.

**FIGURE 1.**
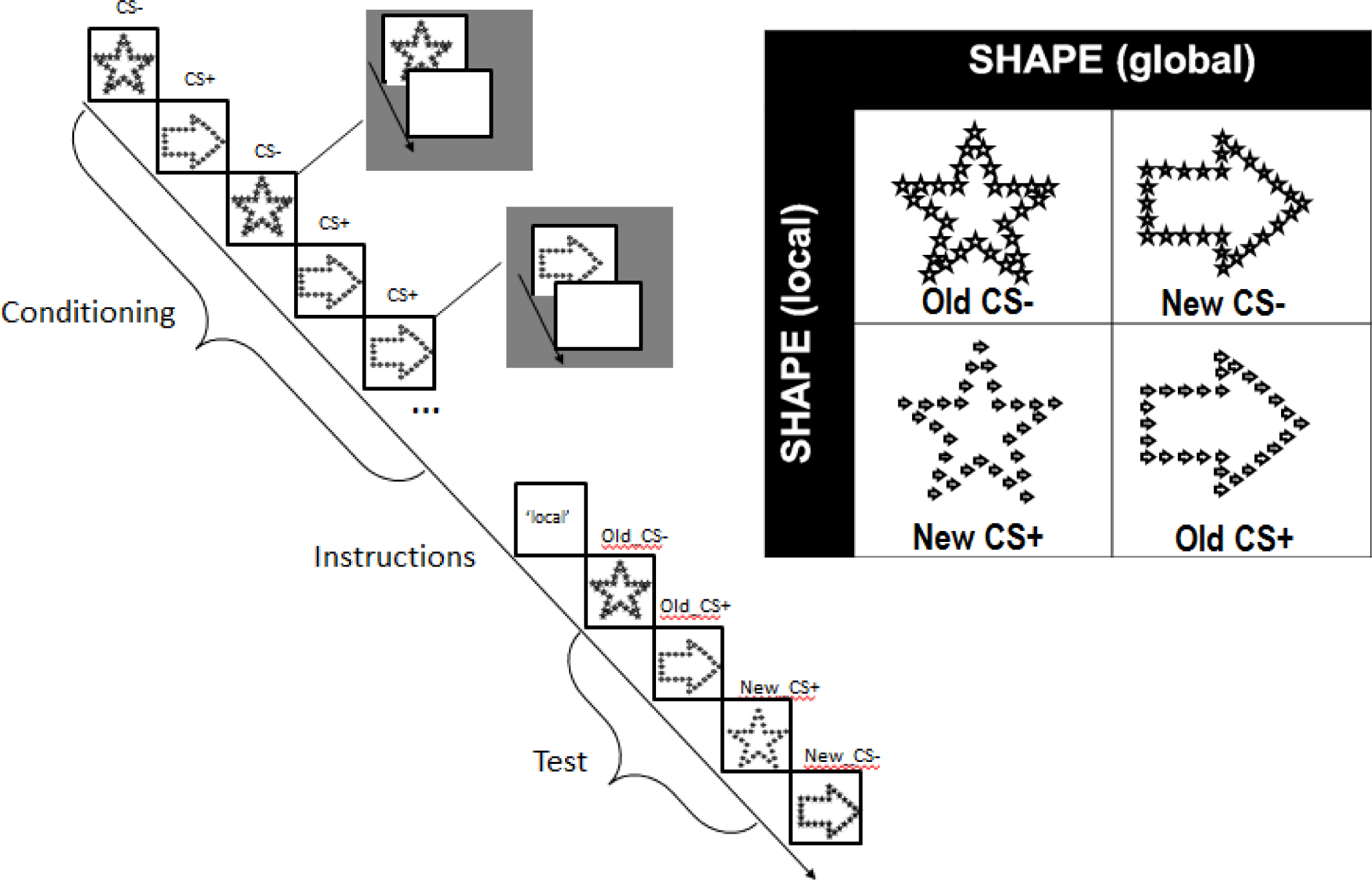
EXPERIMENTAL TASK. **Left:** timeline of a single block in the experimental task, including the conditioning, instructions, and test stages. **Right:** CSs in this block were drawn from the 4-figure subset crossing the local and global dimensions of the star and arrow shapes.

#### US

The majority of studies of VMAC use rewarding USs, but there is also evidence for VMAC with aversive outcomes, including pain (e.g. Wang et al., 2013). Here the US was a painful electric stimulation delivered to the back of the right hand (see Apparatus, above).

### Procedure

On arriving at the laboratory, participants were given an information sheet informing them of the justification for the study and of the use of electrical stimulation. After they signed the consent form, participants were fitted with the electroencephalogram (EEG) cap, and sat in a dimly-lit and sound-attenuated room, 90 cm in front of the monitor screen, where an electrode was attached to the dorsum of their right hand. Once the electrode was attached the participants undertook a series of procedures, described below, in the following order: pre-experiment rating of materials, pain calibration, habituation, experimental task, and post-experiment rating of materials.

#### Pre- and post-experiment rating of likability and contingency

Participants were presented with all of the figures and rated how much they liked each one using a 5-point Likert scale (likability rating task). They then saw all figures again, and guessed, by entering a percentage, how likely each figure was to be followed by a painful stimulation (contingency rating task). The order of the figures in each rating task was randomised for each participant. The likability and contingency rating tasks were repeated at the end of the experiment.

#### Pain calibration

This procedure ensured that participant could tolerate the stimulation, and that the stimulations were psychologically equivalent across participants. During this procedure participants received a series of stimulations starting from 0.4mA, and incrementing by 0.4mA at each step. Participants rated each stimulation on a scale from 0 – 10 where a score of 0 reflected not being able to feel the stimulation, 3 reflected a stimulation level that was on the threshold of being painful, 7 related to a stimulation that was deemed painful, but still tolerable, and 10 related to ‘unbearable pain’. The scaling procedure was terminated once the participant reported the level of stimulation as being equivalent to ‘7’ on the scale. This calibration procedure was performed twice to allow for initial habituation/sensitisation to the stimulation. The power levels that induced a rating of ‘7’ on the second run of the calibration procedure were used as US. The US was therefore a painful but tolerable sensation.

#### Habituation and method of CS presentation

Participants passively viewed a randomised list of all of the CS figures. CS figures were displayed at the centre of the screen, a 17’’ monitor with a resolution of 1024×768 pixels and a refresh rate of 60Hz. The duration of the presentation of each CS was 3,300ms. That time included 66 on-off cycles in which CS figures were displayed on a uniform white background for 33.3ms (‘on’) and the screen turned black for 16.6ms (‘off’), resulting in a 20Hz flickering display. The inter-trial interval between CSs was 2,500ms, during which the screen was white.

#### Experimental task

We designed a novel task to reveal model-based Pavlovian control of VMAC. A schematic of the task is shown in Figure 1. The task progressed through two stages - a conditioning stage with 24 trials and a test stage with 4 trials, which are described in detail below. Each trial included the presentation of a CS; when this was a CS+, the trial always terminated with US delivery. Crucially, the logic of the task necessitated an extremely brief test stage that yielded only a single trial for the contrast of interest. This was necessary in order to ensure that VMAC could not be controlled through a model-free process; once new_CSs were reinforced, that reinforcement could inform the value assigned to new_CSs in their second presentation. Therefore, we needed to measure attention to new_CSs upon their first presentation, before they were reinforced, to prevent any possibility that threat value could be informed by the experience of reinforcement. This requirement led us to measure attention using SSVEPs (Matthias M. Müller et al., 1998). SSVEPs have excellent SNR compared to ERP and time-frequency analysis of EEG measurements (Norcia et al., 2015; Wieser et al., 2016), which can even allow a measure of effects at the single-trial level (Keil et al., 2008). This is because the neurons that generate the SSVEP signal oscillate at the driving frequency, which is precise and known a-priori, such that it is less affected by background EEG noise. Even within the narrow band, the time-locking of oscillation phase to the stimulus (here, the CS) decreases the impact of non-experimentally-driven oscillations. In addition, a-priori knowledge of the frequency band and the electrodes sensitive to it prevents the need to correct for multiple comparisons.

The task was repeated once in each of 24 task blocks. Each task block used novel stimuli, as described in the material section, preventing the transfer of learning across blocks. Each task block lasted 2.5 minutes, with a 5-second break between blocks. Participants practiced the experimental task before it commenced in two practice blocks, using the practice materials described above.

Before the experimental task began participants were given instructions for the experimental task. They were asked to fixate on the fixation cross throughout each block, to observe the figures, and to pay attention to the relationships between the figures and the pain. To encourage compliance, participants were told that their memory for these associations will be tested. This instruction does not privilege memory for the CS+ compared to the CS−, and therefore cannot be responsible for observed threat responses.

##### Conditioning stage

During the conditioning stage, participants learned that one figure (old_CS+) always predicted pain but another (old_CS−) was safe. CSs were fully predictive of their respective outcomes to reduce any effects of stimulus predictability and of uncertainty, which are tightly intertwined with the effect of value on attention control (Le Pelley et al., 2016). We used previous trial-by-trial dissection of threat effects on the SSVEP signal (Wieser, Miskovic, Rausch, & Keil, 2014) to decide how many conditioning trials were necessary in the conditioning stage. They observed a significant modulation of the SSVEP by aversive reinforcement was observed after 5-10 conditioning trials. Therefore, here we used 12 conditioning trials with each CS. The old_CS+ figure and the old_CS− figure were presented 12 times each, at a random order. The details of how each CS was presented was the same as during habituation, but here, when old_CS+ figures were presented, the US was delivered during the very last cycle, at the offset of the last ‘on’ screen.

##### Test stage

After the conditioning stage was completed, participants viewed one of three possible instructions for 10s. In the experimental condition the instruction was the word ‘global’ or the word ‘local’. These words indicated the terms under which the US was to be delivered in the test stage, namely, whether the global or local attribute of the old_CS+ would be reinforced. In the control condition the instruction was a meaningless alphanumeric string, which gave participants no information as to which attribute of the old_CS+ would be reinforced.

Four trials were presented after the instructions. The first two included the presentation of old_Css (their order was randomised), and the last two the presentation of new_CSs (their order was also randomised). New_CSs were the “other” two figures from the same four-figure subset from which the old_CSs were drawn. As can be appreciated from examining Figure 1, each of the new_CSs consisted of one previously-reinforced attribute and one previously-safe attribute. The old_CS+ and the new_CS+ were reinforced; the old_CS− and the new_CS− were not.

Participants did not see the new_CSs before, so without the instructions they could not predict which one would be reinforced. The only way for participants to predict whether the US will follow the new_CS+ or the new_CS− was to infer this from the instructions by drawing on their memory of old_CSs. For example, after the instruction ‘local’ participants who remembered the local attribute of the old_CS+ could infer that (1) the global attribute of new_CSs did not determine whether the US will be delivered or not (2) the US will follow any new_CS with the same local attribute as the old_CS+. Taken together, such participants would predict pain after the new_CS+ but not after the new_CS−. New_CSs were reinforced in accordance with the instructions, confirming participants’ expectations. Old_Css were reinforced in accordance both with the contingencies established during the conditioning stage and the instructions.

While in previous research a newly-acquired conditioned response could be observed within 5-10 trials with each CS (Wieser et al., 2014), the test stage in each task block here only yielded just one trial in each condition. To increase SNR the same structure described thus far – a conditioning stage followed by the test stage - was repeated in each task block. 16 task blocks were allocated to the experimental condition (8 with ‘global’ and 8 with ‘local’ instructions) and 8 were allocated to the control condition.

### EEG recording and analysis

#### EEG recording

Continuous EEG recordings were obtained from a 64-channel cap with in-build electrodes (Biosemi Active Two) using the 10-20 configuration system. Data were digitised at a rate of 2048Hz and filtered online between 0.1 and 100 Hz. The recording was referenced online to the Common Mode Sense active electrode. The Driven Right Leg passive electrode was used as ground. The impedance was kept below 40kΩ. Eye movement and blinks were recorded from horizontal and vertical electro-oculogram.

#### Preprocessing of EEG data

Data were analysed using SPM12. They were converted from their native format and then filtered with three 2 order Butterworth IIR zero-phase forward and reverse filters: a 1Hz highpass, a 80Hz lowpass, and a 49.50Hz-50.5Hz notch filter to remove mainline noise. Data were downsampled to 200Hz and re-referenced to the average of all electrodes. Eye blinks and saccades were marked on the VEOG and HEOG channels (or Fp1 in two participants) using an automatic algorithm that was thresholded separately for each participant.

Further pre-processing was conducted for the purpose of complex demodulation (see below). Data were segmented between −600ms before the onset of CSs to 3250ms after CS onset (Just before the offset of the CS/US onset, 3300ms after CS onset). Segments where the following artefacts were present on occipital channels (Oz, POz, O1, O2, O3, O4) were rejected: jumps greater than 150 µV; peak-to-peak differences greater than 250 µV; flat segments. Channels where more than 20% of the trials were rejected were excluded from analysis. This left, on average, 282.62 learning trials and 7.83 test trials with each CS in each condition. Artefacts associated with eye blinks and saccades were corrected using the singular value decomposition (SVD) technique implemented in SPM12 which captures eye artefacts and removes the associated component.

#### Complex demodulation

Threat effects were operationalised as an increased response to the CS+ compared to CS−. Previous work suggested that threat effect are more pronounced later in the presentation of the CS, because the threat response is greater when threat is imminent, and the perceptual processing of predictive sensory features of the CS is enhanced only when the US is imminent (Miskovic & Keil, 2012; Paterson & Neufeld, 1987). Complex demodulation was therefore carried out to determine exactly when threat effects were present. Complex demodulation was conducted using SPM12 on the entire segment, with the multitaper method, a hanning window, and a resolution of 1Hz. Data in each condition were averaged using robust averaging, a method that down-weights outliers (Litvak et al., 2011). The signal from electrodes Oz and POz was extracted around the driving frequency (19-21Hz). These data were averaged across all 12 trials in each condition and the 24 blocks of the experimental task (288 trials for each CS). In agreement with Miskovic and Keil (2012), threat effects were greatest during the second half of the presentation of the CS (Figure 2). An examination of the topographies associated with threat supported our selection of electrodes of analysis. We used these results to constrain our spectral analysis.

**FIGURE 2.**
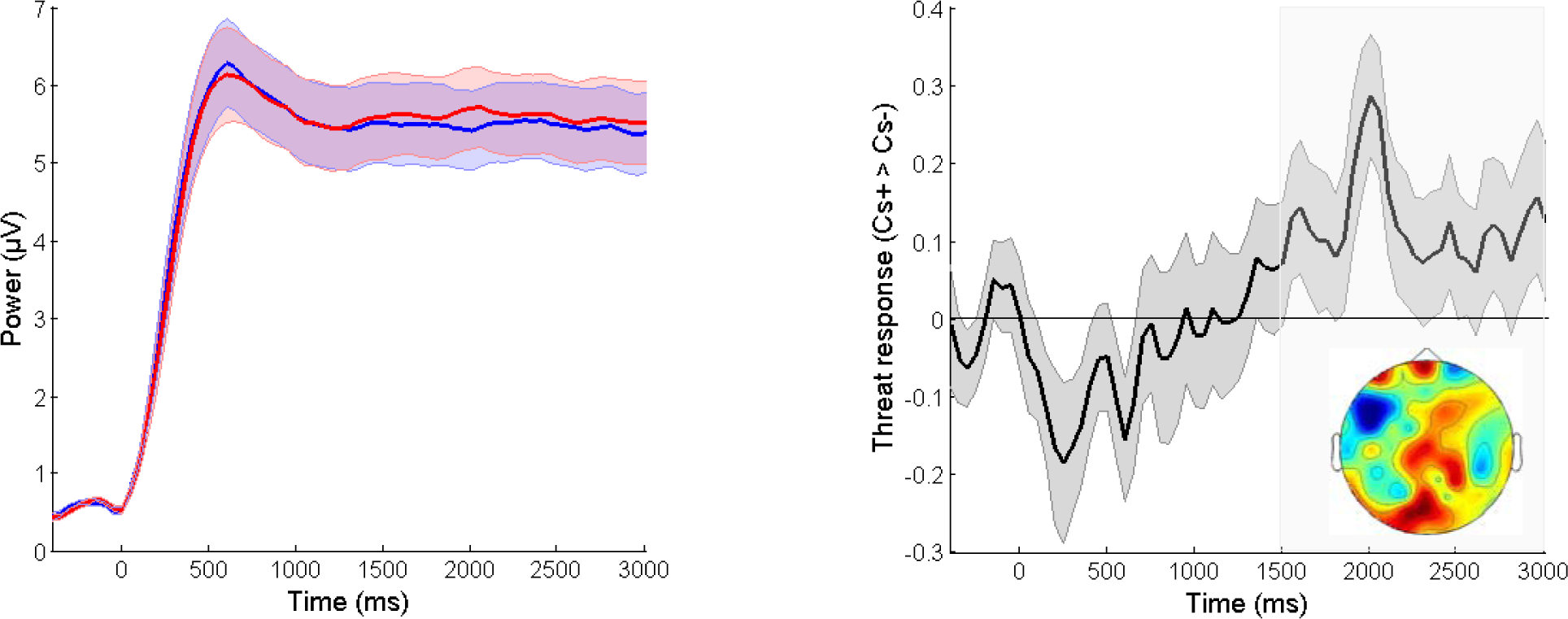
THREAT EFFECTS IN THE CONDITIONING STAGE AS A FUNCTION OF TIME FROM CS ONSET. **Left.** SSVEP signal amplitudes in the conditioning stage for CS+ and CS−, extracted from occipital electrodes Oz and POz, and averaged across 19-21Hz. Shaded areas plot the standard error. **Right.** Threat effects in occipital electrodes Oz and POz, operationalised as the difference between CS+ > CS−, are plotted as a function of time from CS onset, averaged across 19-21Hz. The time window 1500-300ms (shaded grey) was used in all analysis of threat effects. The topology inset shows that the threat effects across that time window.

#### Spectral analysis

Based on the results of the complex demodulation step, spectral analysis was conducted using the spatiotemporal window of 1500-3000ms from CS onset, at Oz and POz. We followed the method in Keil (2008) to maximise sensitivity to differences of interest by minimising 20Hz signal that does not keep in phase with the stimulation. Within the time and spatial windows defined above, 26 windows of 250ms (5 cycles of the SSVEP) were segmented for each trial. The first window started at 1500ms from CS onset and each subsequent window started 50ms (1 SSVEP cycle) later; the last window started 2750ms from CS onset. For each trial, these 26 windows were then averaged in time, resulting in averages that corresponded to single trials. We then averaged across all of the single trial averages from the same condition (Keil et al. did not perform this last step because they were interested in single-trial data). These condition-wise averages were Fourier-transformed using the FFT function in Matlab.

#### Statistical analysis

Across participants, some of the data significantly diverged from normality, according to the Shapiro-Wilk test, and all data were therefore log-transformed. Data from the conditioning stage in each experimental block were binned (4 bins; 3 trials per bin), averaged across the 24 task blocks, and analysed with a 2 (threat: CS+, CS−) × 4 (bins in the conditioning stage). The main hypothesis concerned the response to new_CSs during the test stage. The response to new_CS+ was compared to the response to new_CS− across the 16 blocks in the experimental condition (collapsing across the 8 blocks in the ‘global’ and the 8 blocks in the ‘local’ conditions) using a one-sample t-test. These responses were also compared to the response to responses to the two ambiguous CSs that followed the 8 control blocks where no instructions were given using two Bonferroni-corrected two-sample t-tests (the averages that went into these comparisons comprised of 16 trials in each condition).

## Results

### Behavioural results

#### Sample selection

Behavioural and EEG data were first checked to verify the presence of a conditioned threat response. We excluded participants who may have disengaged from the task based on their ‘learning score’, computed as the increased contingency ratings given to old_CS+ stimuli after the experiment compared to the pre-experiment rating given to the same stimuli. Increased ratings must be based on learning that occurred during the experiment, and therefore reflects at least a minimal level of engagement. The learning score purposefully ignores the magnitude of the conditioned response, which could be computed as differential ratings of the old_CS+ and old_CS−, in order not to bias the selection of the sample. To be included participants had to show a numerical (above 0) increase in ratings. Based on this criterion, three participants were excluded from analysis, leaving a sample of N=20.

#### Manipulation check

Contingency and liking ratings for the stimuli were entered into two separate 3-way repeated-measures ANOVAs with the factors time (pre-experiment, post-experiment), threat (CS+, CS−), and status (old, new). The results evidenced a conditioning effect (Figure 3). In both the analysis of contingency ratings and in the (separate) analysis of liking ratings the 3-way interaction between time (pre-experiment, post-experiment), threat (CS+, CS−), and status (old, new) was significant (contingency: F(1,19)=34.95, P<.001, partial eta=.65; liking F(1,19)=4.79, p=.04, partial eta=.20). We unpacked this interaction by examining the old and new CSs separately. The 2-way interaction was significant for old_CSs (contingency: F(1,19)=59.98, p<.001, partial eta=.76; liking: F(1,19)=17.89, p<.001, partial eta=.48) as well as for new_CSs (contingency: F(1,19)=4.65, p=.04, partial eta =.20; liking: F(1,19)=9.16, p=.007, partial eta=.32). The results showed that participants expected the US more frequently following the CS+ and liked the CS+ less than the CS−, and that these effects were stronger for old_CSs than for new_CSs, probably because old_CSs were repeated multiple times, but new_CSs were only presented once.

**FIGURE 3.**
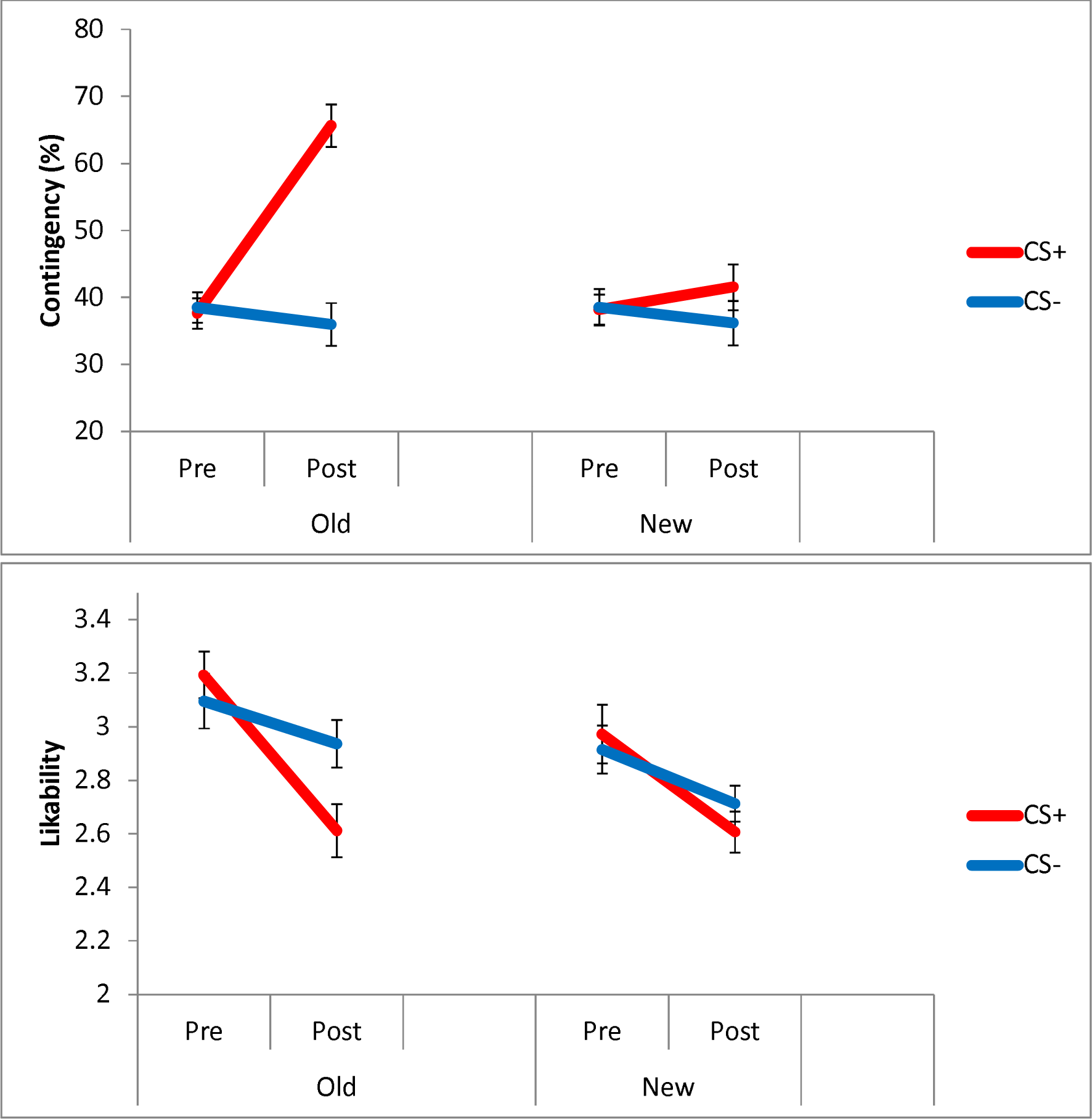
CONTINGENCY AND LILKABILITY OF CONDITIONED STIMULI. The contingency and likability ratings of stimuli used as CSs before and after the experimental task. Error bars indicate the standard error of the mean.

### EEG results

#### Manipulation check

Threat effects on the SSVEP during the conditioning stage were analysed with a 2 (threat) × 4 (trial bins) repeated-measures ANOVA. The successful manipulation of threat in this experiment was evident in differential responses to the CS+ and CS− during the conditioning stage (Figure 4), F(1,19)=5.50, p=0.02, partial η^2^ =0.22. Although numerically the magnitude of threat effects was only observable after 3 trials, the interaction of threat and trial bins was not significant, F(3,57)<1, suggesting that they remained consistent across each of the experimental blocks. The main effect of binned trials, F(3,57)=6.98, p<.001, partial η^2^ =0.27, was also significant, denoting an overall decrease in attention as the block progressed. To verify that threat responses were also obtained in the test stage, responses to old_CSs in the control condition, where participants had no reason to make any model-based inferences, were analysed with a one-tailed paired t-test, contrasting old_CS+ and old_CS. We observed differential responses to the old_CS+ and old_CS−, t(19)=2.36, p=.015 (one-tailed), Cohen’s d = 0.53. Because only the data from the conditioning stage were pre-selected, the data from the test stage provide a useful confirmation.

**FIGURE 4.**
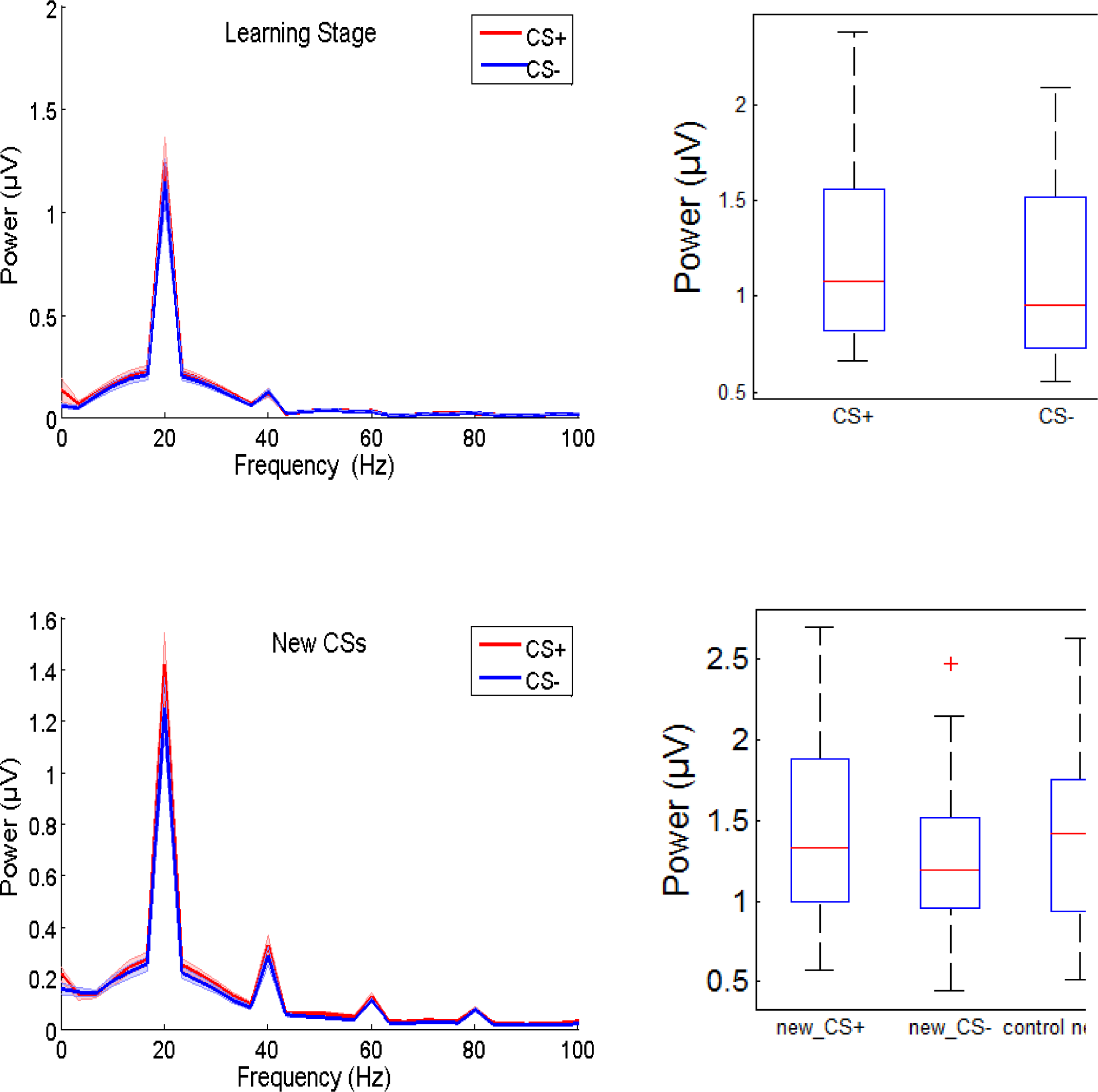
SPECTRAL ANALYSIS OF THREAT EFFECTS. **Left.** Spectral analysis of signal in the conditioning stage for CS+ and CS−, extracted from occipital electrodes Oz and POz at the 1500-3000ms time window, showing that threat modulated the 20Hz SSVEP signal and some of its harmonics. **Right.** The magnitude of the 20Hz threat effect, showing the variability of this effect across participants. The red line indicated the mean; the box indicates the inter-quartile range.

#### Main hypothesis

Our main hypothesis was that responses to the new_CS+ would be greater than responses to the new_CS−. The hypothesis was evaluated with a one-tailed paired t-test comparing SSVEPs to the new_CS+ and new_CS−. As predicted, SSVEP amplitudes were higher during the presentation of the new_CS+ compared to the new_CS−, t(19)=2.22, p=.02, Cohen’s d = 0.50. Additional 2-tailed t-tests, controlled for multiple comparisons with a p-value of 0.025, compared each of these SSVEPS to the SSVEP elicited by ambiguous new_CSs in the control condition. SSVEP amplitudes were equivalent during the presentation of the new_CS+ and the new_CSs in the control condition, t<1, suggesting that participants experienced ambiguous figures as threatening when they could not use the instructions to disambiguate them. This interpretation is supported by a significant difference between the new_CSs in the control condition and the new_CS−, t(19)=2.58, p=0.018, Cohen’s d = 0.58, suggesting that VMAC was attenuated when participants knew that pain was unlikely.

## Discussion

During the learning stage of our Pavlovian conditioning task, we observed an increase in the amplitude of the SSVEP signal towards stimuli with learned aversive value. These results are not surprising, given much evidence that the valuation system can control attention allocation (Le Pelley et al., 2016), but important because they confirm relatively limited evidence for Pavlovian control of VMAC towards stimuli with aversive value (Van Damme et al., 2006; L. Wang et al., 2013; Wentura et al., 2014; Wieser et al., 2014), which is less established than Pavlovian control of VMAC towards reward, or the effect of value on instrumental control of VMAC. Uniquely, these results also suggest a way to observe the neural evolution of VMAC across the learning process, and even in the first trial, something that has not been possible using other neuroimaging investigation (Olsson & Phelps, 2007; Phelps et al., 2001) or in animal models (Balcarras, Ardid, Kaping, Everling, & Womelsdorf, 2016).

Our key result was that SSVEP amplitudes were larger when participants were presented with a new shape that they inferred predicted physical pain (the new_CS+), compared to a new shape that they inferred predicted safety (the new_CS−). Because the amplitude of SSVEPs is higher for attended stimuli compared to unattended ones (Matthias M. Müller et al., 1998; Muller et al., 1998), our findings suggest that more attention was allocated to the new_CS+ compared to the new_CS−. We argue that the differential attentional response to new_CS+ and new_CS− suggests a Pavlovian model-based control of VMAC, an argument that we dissect in the ‘theoretical considerations’ section, below. It has to be noted that while the SSVEP is known to be sensitive to VMAC such as in aversive learning (Kastner-Dorn, Andreatta, Pauli, & Wieser, 2018; Miskovic & Keil, 2013; Wieser et al., 2016) and to emotional stimuli in general (Keil et al., 2003; Keil, Moratti, Sabatinelli, Bradley, & Lang, 2005; McTeague, Shumen, Wieser, Lang, & Keil, 2011; Wieser et al., 2016), heightened ssVEP amplitudes are also found in response to increased working-memory load (Silberstein, Nunez, Pipingas, Harris, & Danieli, 2001), and for attended relative to unattended stimuli (Morgan, Hansen, & Hillyard, 1996) (Hillyard et al., 1997)(M. M. Müller, Malinowski, Gruber, & Hillyard, 2003; Matthias M. Müller et al., 1998). Thus, the enhanced SSVEP amplitudes for may reflect any of these respective processes.

By contrast to the experimental blocks, where the value of new_CSs could be inferred through a combination of the instructions and stored memories of the learning stage, in control blocks no instructions were given. Therefore, the threat value of control new_CSs was ambiguous, and participants could not predict which one would be followed by pain. These ambiguous new_CSs in the control condition attracted increased attention compared to the new_CS−. This result, which suggests orienting towards ambiguous stimuli, accords with previous findings, where instructed, ambiguous, novel CSs gave rise to increased physiological arousal and increased activation in the amygdala and the insula (Phelps et al., 2001).

Our study confirms other demonstrations where instructions about Pavlovian contingencies encourage responses that mimic the effect of associative learning through experience (reviewed in Mitchell, De Houwer, & Lovibond, 2009). This is the first demonstration that propositional information – a form of instruction – influences Pavlovian control of attention allocation. This demonstration is particularly important because a previous experiment suggested that in Pavlovian tasks attention allocation obeys associative learning principles, and is immune to propositional knowledge. In Moratti and Keil’s study (Moratti & Keil, 2009) the SSVEF during CS presentation (steady-state visual evoked field, measured with MEG) increased with increased number of sequentially reinforced CSs, not with increased US expectancy, which, in turn, only increased when previous CSs have consistently not been reinforced. Indeed, other studies have also observed that similar ‘gambler’s fallacy’-like paradigms give rise to conditioned responses that are based on model-free, not model-based value (Clark, Manns, & Squire, 2001; Perruchet, 1985). The results of Moratti and Keil, indicative of attention allocation, were particularly intriguing because they appeared to contradict evidence that expectancy influences visual attention (Downing, 1988). Taken together, it appears that associative mechanisms dominate attention allocation in the gambler’s fallacy paradigm, at least when using a delayed conditioning procedure (Clark et al., 2001), while in other paradigms – including those using delayed conditioning, as we did here – propositional information holds more sway. It is possible that the balance between these mechanisms is affected by the certainty of each system in its threat evaluation (Daw et al., 2005).

While attentional responses were affected by propositional information here, it is possible that other classically-conditioned responses were not. In particular, because attention allocation was affected in the first trial it bears stronger resemblance to US-expectancy ratings and to classically-conditioned skin conductance responses, which were immediately influenced by instructed extinction, than to potentiated startle, which was not (Sevenster et al., 2012). Further research is required to examine this potential dissociation using our paradigm, and to verify whether instructed extinction, like instructed threat, also alters VMAC instantaneously.

By using neural measures to index control of VMAC we move a little closer to understanding how propositional information is implemented at the level of the neurobiological mechanism. Increased SSVEPs during the presentation of threatening stimuli is thought to be driven by re-entrant connections from the amygdala, ACC and OFC, which amplify the processing of adaptive information (Miskovic & Keil, 2012). Repeated pairing between a stimulus and pain can change the neural representation of the pain-predicting stimulus. For example, repeated pairing between a tone and a painful shock change the tuning frequency of neurons that encode these tones, and stimulation of the amygdala is sufficient to produce this effect (Chavez, McGaugh, & Weinberger, 2013). Here, however, such a process could not occur because attention modulation was manifested before the reinforcement itself. A meta-analysis of studies of instructed fear found that the dorsomedial prefrontal cortex is uniquely associated with a conscious appraisal process (Mechias, Etkin, & Kalisch, 2010). Similarly, the same region has been shown to dynamically modulate model-free valuation in the OFC, striatum, and hippocampus (Li, Delgado, & Phelps, 2011). It is therefore likely that increased response to the new_CS+ was due to projections from the dorsomedial prefrontal cortex to the OFC and ACC, regions that are strongly connected to the amygdala and able to modulate its activity (Lee, Heller, van Reekum, Nelson, & Davidson, 2012; Schiller & Delgado, 2010), with downstream re-entrant effects in the visual cortex.

The constrained data yield of the paradigm should be acknowledged as a limitation of this study. While the effect sizes in all of the statistical tests were all of a ‘medium’ size, according to Cohen’s classification (Cohen, 1988), the study should be replicated in order to increase confidence in this novel result. For the same reason, we could not explore the influence of ‘dimension’ (global or local) in the results we obtained, because this would have halved the number of trials that we could analyse.

### Theoretical considerations

We argue that increased SSVEPs to new_CS+ compared to new_CS− in this experiment was likely due to a Pavlovian, not an instrumental process The paradigm was entirely passive; pain outcomes were independent of participants’ behaviour or how they allocated their attention. Participants could not benefit from allocating differential attention to specific CSs. Indeed, participants were told explicitly, and also knew through experience across the 24 blocks of the task, that the stimulation levels were pre-determined and that they could therefore not influence it. Participants had no reason to allocate more attention to the new_CS+ in order to increase success in the post-experiment rating task, because they have already completed it once before they started the experiment, and knew, therefore, that performance would benefit equally from attending all of the stimuli.

Although the control of VMAC here could not influence objective outcomes, it is possible, in principle, that it incurred some internal benefit. Specifically, paying extra attention to threatening CSs here may have decreased subjective pain. It is important to consider this possibility because expected and experienced pain are not true reflections of objective tissue damage. Instead, pain experience is strongly modulated by pain expectations (Atlas & Wager, 2012; Berns et al., 2006; Morley, Vlaeyen, & Schrooten, 2012; Paterson & Neufeld, 1987; Tabor, Thacker, Moseley, & Körding, 2017; Vlaev, Seymour, Dolan, & Chater, 2009), which modulate endogenous analgesic mechanisms (Anchisi & Zanon, 2015; Tracey, 2010; Wager et al., 2004), and experimental pain expectations are themselves influenced by pre-existing individual biases (Hoskin et al., n.d.).

Yet closer scrutiny suggests that it is unlikely that attending the new_CS+ triggered endogenous analgesia. While participants needed to attend experimental new_CSs to decipher exactly which one predicted pain and which one did not, this need not result in differential attention to the two new_CSs. The US was completely predictable, always of the same intensity, and presented at the same time, so attending the new_CS+ is unlikely to have altered pain expectations meaningfully. It is possible that attending the new_CS+ increased the precision of expectations (Kok, Rahnev, Jehee, Lau, & de Lange, 2012). However, expecting high pain with greater certainty would increase subjective pain, not decrease it (Hird et al., 2018). In fact, much evidence suggests that distraction, not attention, is an effective pain-coping strategy (Buhle, Stevens, Friedman, & Wager, 2012; Eccleston, 1995; Sharar et al., 2016; Weiss, Dahlquist, & Wohlheiter, 2011).

To fully establish that a response is controlled by a Pavlovian process researchers have utilised omission schedules, where the response incurs a cost (Le Pelley et al., 2016; Mackintosh, 1983). This method is achievable for model-free Pavlovian responses, but in the case of model-based Pavlovian control known eventual costs should, by definition, alter the world-model that inspired the responses in the first place. It is therefore potentially tricky to utilise this method to test that control was Pavlovian and model-based. In summary, although the feeling of pain is malleable, and we have not conclusively demonstrated that attention was controlled through a Pavlovian rather than an instrumental process, we can presently think of no a-priori reason that attending the new_CS+ in this experiment would be advantageous. The intuitive sense that we would all want to look – that we might not be able to attend anything else – is perhaps simply the reflection of Pavlovian misbehaviour (Dayan et al., 2006).

During the test stage participants could combine the propositional information provided to them in the instructions (about the dimension that will be reinforced - global or local) with their stored representation of the reinforced attribute (the particular shape that predicted the US during the conditioning stage) to form a prediction of the value of new_CSs. That they have, indeed, done so is evident in the differential attention they allocated to new_CSs in experimental blocks. While we did not test the model-based nature of control of VMAC formally, e.g. by using a two-step task (Otto, Skatova, Madlon-Kay, & Daw, 2015), increased attention to the new_CS+ compared to the new_CS− here involved “prospective cognition, formulating and pursuing explicit possible future scenarios based on internal representations of stimuli, situations and environmental circumstances” – the hallmark of model-based Pavlovian control according to Dayan and Berridge (2014, p. 5).

While new_CSs were constructed such that previous experience would render them equally ‘threatening’ and ‘safe’, and render it difficult for a model-free algorithm to implement differential responses to these CSs, it is possible that the instructions created a model which then trained a model-free controller, as in schemes such as Dyna (Sutton, Szepesvári, Geramifard, & Bowling, 2008) or “biased” learning (Doll, Jacobs, Sanfey, & Frank, 2009), or, alternatively, trigger an existing model-free controller. Regarding the first alternative, it is clear that the limits of model-free reinforcement learning are constantly expanding with the introduction of meta-reinforcement learning (J. X. Wang et al., 2016). Future computational work could explore whether indirect reinforcement learning can link stored knowledge and propositional information within a few seconds and a single trial to influence the value predicted in a novel state. In the “biased” learning scheme, for example, the training of the model-free controller relied on a modulation of the value of reinforcers (Doll et al., 2009), so it may not be realistic to expect such training in measurements that take place prior to any reinforcement, as in the present paradigm.

The second alternative is that the instructions created a model that triggered model-free habits to control VMAC. Organisms may habitually attend more highly valued (compared to neutral) stimuli preferentially because this incurs reward over the long-term, even though it does not do so in a particular scenario. Attention to valued stimuli could improve the encoding of CSs, strengthen their memory traces, and thus facilitate optimal decisions when the opportunity arises to act on the same stimuli (Lieder, Griffiths, & Hsu, 2018). Additionally, prediction error minimisation – something that is considered globally adaptive (Pezzulo, Rigoli, & Friston, 2015) - may be facilitated if the excellent encoding of previously-valued stimuli increases the precision of the model we have of the world around us. The model constructed by the instructions could therefore simply indicate to the system which stimuli are likely to have large absolute value, as well as which have value which is highly uncertain; but the actual attentional control may be carried out through the habitual mechanism.

Finally, it is possible that VMAC was, indeed, controlled by a model-based, Pavlovian process. In that case, following the experimental instructions, participants may have constructed a model of the new_CSs and their predictive value by combining the new propositional information and stored internal representations. It is possible that while they viewed the instructions, participants recalled old_CSs and generalised their aversive value to imagined new_CSs. It is also possible that participants volitionally inhibited the representation of the global or local dimensions of memorised learned CSs as well as actual new_CSs, to support generalisation from the learning to the test stage. Dayan and Berridge (2014) discuss such recall and revaluation processes as mechanisms that allow model-based Pavlovian control.

At the experiential level, increased attention to the new_CS+ suggests that the information given to participants worked as an emotion regulation technique – it rendered ambiguous stimuli instantly threatening. Drawing the connection between model-based and model-free control, on the one hand, and cognitive and emotional control, on the other (Sevenster et al., 2012) can help the quest to ground emotion regulation and behaviour change techniques more tightly in computational theories (Etkin, Büchel, & Gross, 2016).

## Acknowledgements

We thank N. Chater and C. Frith for the discussions that inspired this paradigm, A. Wilkinson for help with data collection, J. Taylor and C. Charalambous for help with data analysis, and M. Bauer and L. Hunter for helpful comments. DT acknowledges the support of ESRC First Grant ES/I010424/1.

